# A Comparative Machine Learning Study of Connectivity-Based Biomarkers of Schizophrenia

**DOI:** 10.1101/2024.01.05.573898

**Authors:** Victoria Shevchenko, R. Austin Benn, Robert Scholz, Wei Wei, Carla Pallavicini, Ulysse Klatzmann, Francesco Alberti, Theodore D. Satterthwaite, Demian Wassermann, Pierre-Louis Bazin, Daniel S. Margulies

## Abstract

Functional connectivity holds promise as a biomarker of psychiatric disorders. Yet, its high dimensionality, combined with small sample sizes in clinical research, increases the risk of overfitting when the aim is prediction. Recently, low-dimensional representations of the connectome such as macroscale cortical gradients and gradient dispersion have been proposed, with studies noting consistent gradient and dispersion differences in psychiatric conditions. However, it is unknown which of these derived measures has the highest predictive capacity and how they compare to raw connectivity. Our study evaluates which connectome features — functional connectivity, gradients, or gradient dispersion — best identify schizophrenia. Figure 1 summarizes this work.

Surprisingly, our findings indicate that functional connectivity outperforms its low-dimensional derivatives such as cortical gradients and gradient dispersion in identifying schizophrenia. Additionally, we demonstrated that the edges which contribute the most to classification performance are the ones connecting primary sensory regions.

## I. Introduction

Functional connectivity holds promise as a potential biomarker for psychiatric disorders^1,2^, as evidenced by a robust body of literature that highlights distinct functional profiles between patients with schizophrenia and neurotypical individuals. Prior studies have reported lower connectivity across regions, reduced small-worldness of the resting state networks, and lower functional network segregation^3–5^. However, the high dimensionality of connectivity data, combined with small sample sizes in clinical research, poses a risk of overfitting when the aim is prediction^6–8^.

Recently, low-dimensional representations of the connectome such as macroscale cortical gradients^9,10^ and gradient dispersion^11,12^ have been proposed. The gradients are derived from functional connectivity matrices through dimensionality reduction with the aim to maximize the cumulative amount of variance explained by the resulting components. The first component (or the *principal gradient*) reflects the functional hierarchy of the cortex^9,10,13^, spanning from the primary sensory (*unimodal*) regions to higher-order (*transmodal*) regions. It has been demonstrated to be consistent across individuals^14^.

Dong et al.^15^ revealed that the principal gradient is contracted in schizophrenia, i.e., the primary sensory regions were closer to the higher-order regions in terms of their functional profile, as indicated by their principal gradient values. This finding indicates lower functional differentiation between uni- and transmodal regions. Based on the gradient framework, more differences in the cortical functional hierarchy between neurotypical individuals and subjects with neurodevelopmental and psychiatric disorders have been reported^12,14,16–18^.

As an extension of the gradient framework, gradient dispersion further quantifies the density of local connectivity, known as functional modularity. Specifically, higher dispersion would indicate lower functional modularity within the cluster, whereas lower dispersion would mean higher modularity. Thus, gradient dispersion characterizes functional differentiation across the cortical hierarchy. This measure is of interest to us since changes in the functional hierarchy of the cortex are reported to be idiosyncratic to schizophrenia^2,15,19,20^.

Nevertheless, it is unknown if these connectivity derivatives can discriminate between patients with schizophrenia and neurotypical individuals, or if they outperform raw connectivity. Here, we attempt to identify the features with the largest biomarker potential from a large set of features including functional connectivity, gradients, and gradient dispersion. In addition, we explore the impact of the number of features on the choice of classifier. We also seek to address the question germane to neuroscience and computational psychiatry: when one has a limited number of subjects and a disproportionately rich set of independent variables, how does one justify the choice of features? Additionally, we elaborate on putative functional underpinnings of schizophrenia based on the features with the highest predictive potential.

## II. Methods

### Data & Preprocessing

The present study’s sample was derived from three open-source datasets: COBRE^21^, LA5c study from UCLA Consortium for Neuropsychiatric Phenomics^22^, and SRPBS multidisorder MRI dataset^23^. All data were acquired in accordance with the Declaration of Helsinki. The links to the datasets are provided in the data availability statement below.

The initial cohort for our investigation consisted of 996 individuals; subsequently, 40 subjects were excluded from analysis due to substantial motion artifacts (average framewise displacement (FD) > 0.5mm). The analyzed sample comprised 248 patients with schizophrenia (SCZ) and 688 neurotypical controls (NC; see Table 1). The scanning parameters of each dataset are reported in Supplementary Table 1.

**Table 1:** Demographic statistics of the sample. *These datasets are part of a larger dataset, SPRBS-1600. FD: framewise displacement.

|  | COBRE | LA5c | KTT* | KUT* | SWA* | UTO* |
| --- | --- | --- | --- | --- | --- | --- |
| $N_{SCZ}$ (248) | 59 | 45 | 46 | 43 | 19 | 36 |
| $N_{NC}$ (688) | 81 | 102 | 75 | 159 | 101 | 170 |
| Mean age (std) | 37.3 (12.5) | 32.3 (8.8) | 32.2 (10.2) | 37.7 (13.2) | 30.7 (9.5) | 34.9 (16.5) |
| Sex: female (male) | 33 (107) | 59 (88) | 47 (74) | 89 (113) | 19 (101) | 104 (102) |
| Mean FD (std), mm | 0.27 (0.11) | 0.17 (0.09) | 0.11 (0.05) | 0.15 (0.07) | 0.16 (0.08) | 0.13 (0.07) |

Preprocessing of MRI data was done using fMRIPrep 20.2.1^24^ which is based on Nipype 1.5.1^25^ (Supplementary Methods 1). The preprocessed BOLD time series were parcellated with the Schaefer parcellation (1000 parcels, 7 Yeo networks)^26^. Then, we computed a connectivity matrix (Pearson correlation) for each subject.

### Macroscale Cortical Gradients

For each subject, we computed cortical gradients by applying the principal component analysis (PCA) to the Fisher z-transformed and thresholded connectivity matrix (Figure 2A). Hong et al.^14^ showed that PCA, when applied to thresholded connectivity matrices, yields more reliable gradients compared to the other dimensionality reduction techniques frequently featured in the gradient literature^10,27,28^. We thresholded the connectivity matrices by discarding 90% of the lowest correlation values including negative values. We used Procrustes alignment to align the gradients of all subjects^29^. To avoid introducing dataset-specific bias to the alignment, we used the gradients computed from the group connectivity matrix of the Human Connectome Project (HCP)^30^ as reference gradients. Figure 2B displays mean variance explained across all subjects for 200 gradients. On average, the principal gradient accounted for ∼6% of variance of thresholded connectivity matrices. Collectively, 200 gradients accounted for ∼80% of variance.

**Figure 1.**
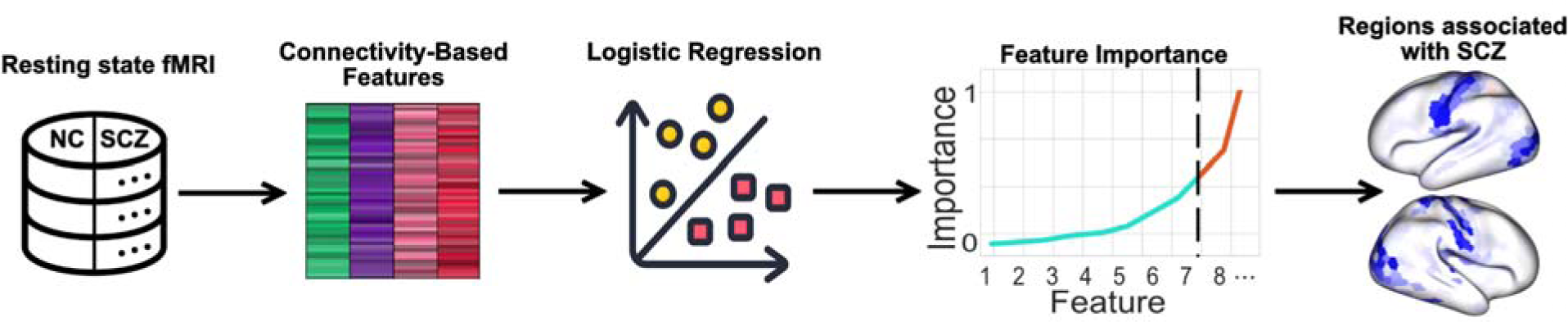
Overview of the methods and main outcome of the paper. Schematic images: Flaticon.com. NC: neurotypical controls, SCZ: patients with schizophrenia.

**Figure 2.**
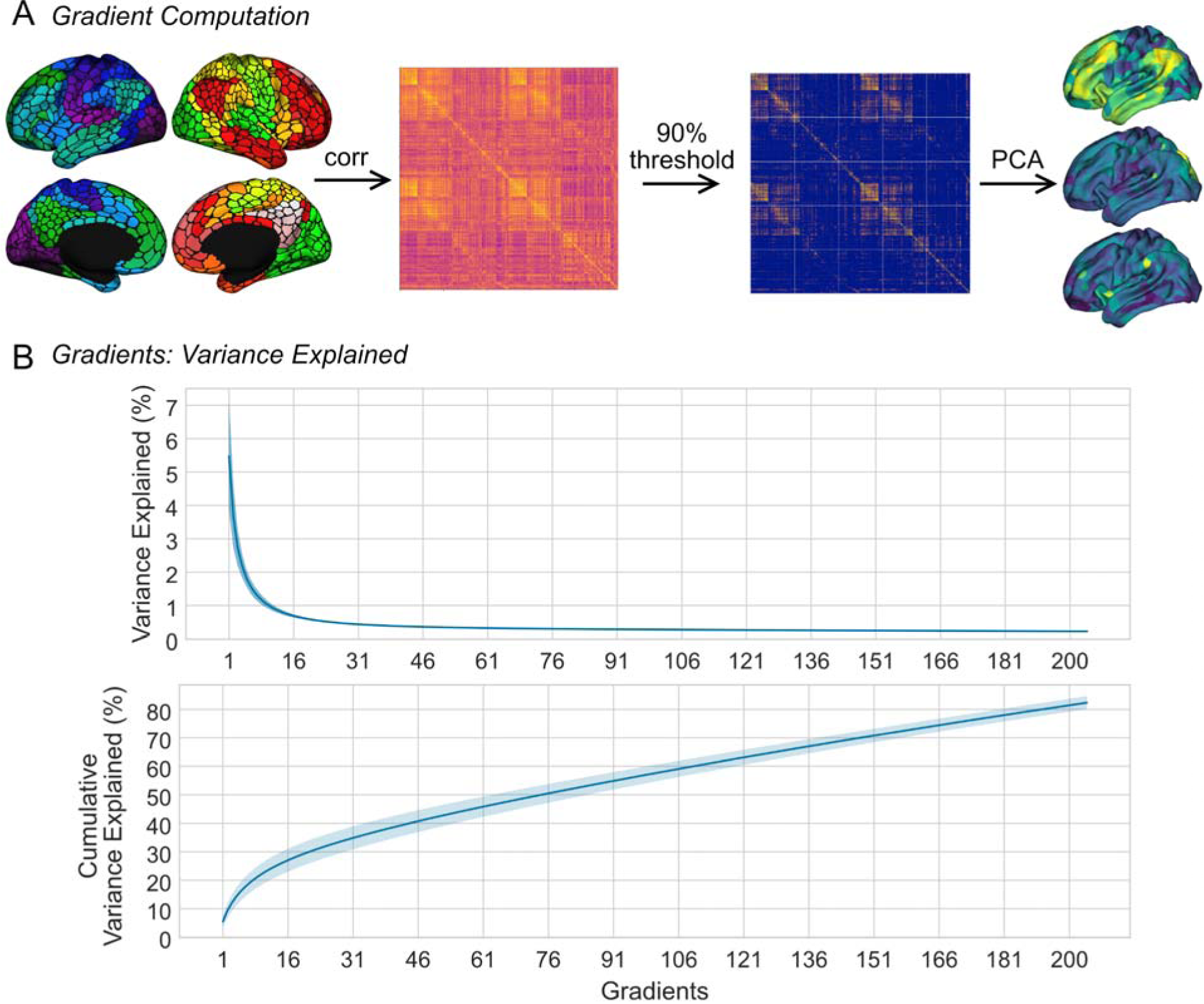
**A:** Parcellated time series (Schaefer atlas, 1000 parcels, 7 Yeo networks^31^) of each subject were correlated to produce a 1000 x 1000 connectivity matrix. Principal component analysis (PCA) was applied to the thresholded matrix to extract 200 gradients. **B:** Variance explained by 200 gradients, mean across subjects ± 1 s.d.

### Centroid Dispersion

Gradient dispersion has been quantified before with different approaches^11,12,32^. These prior investigations, akin to the present study, used dispersion to operationalize functional modularity across the cortex. However, these methods necessitated the identification of centroids of networks for which dispersion was computed relative to the regions encompassed within the network. Thu, dispersion values were only assigned to centroids and categorized as within-or between-network dispersion.

Following this procedure, we computed within- and between-network dispersion for seven Yeo networks^31^ in the 3-dimensional gradient space. Within-network dispersion was quantified as the sum of squares of Euclidean distances from the centroid of the network to the rest of its regions. For the gradient values belonging to a given network, the centroid was defined as three median values of the first three cortical gradients, as in Bethlehem et al.^11^; the position of the centroid in the 3D latent gradient space is defined by these three values. Between-network dispersion was defined as the Euclidean distance between the network centroids. Thus, centroid dispersion amounted to 28 values per subject: 7 values for within-network and 21 values for between-network dispersion. Centroid dispersion is schematically illustrated in Figure 4B (right).

### Neighborhood dispersion

Centroid dispersion can vary depending on how the networks are delineated (e.g., if a different number of networks is used). Hence, we computed neighborhood dispersion with the aim to circumvent this potential confounding factor. Specifically, we calculated dispersion for individual regions which enabled us to maintain the spatial resolution congruent with the gradients (1000 regions per measure).

To compute neighborhood dispersion for every region, we identified K closest neighboring regions (Figure 3) via the K-Nearest Neighbors (KNN) algorithm. Then, we computed the mean Euclidean distance between the focal region and its designated neighboring regions in the gradient space. The resulting value quantified dispersion for that specific region. Neighborhood dispersion was computed for combinations of gradients spanning from 1 to 200.

**Figure 3.**
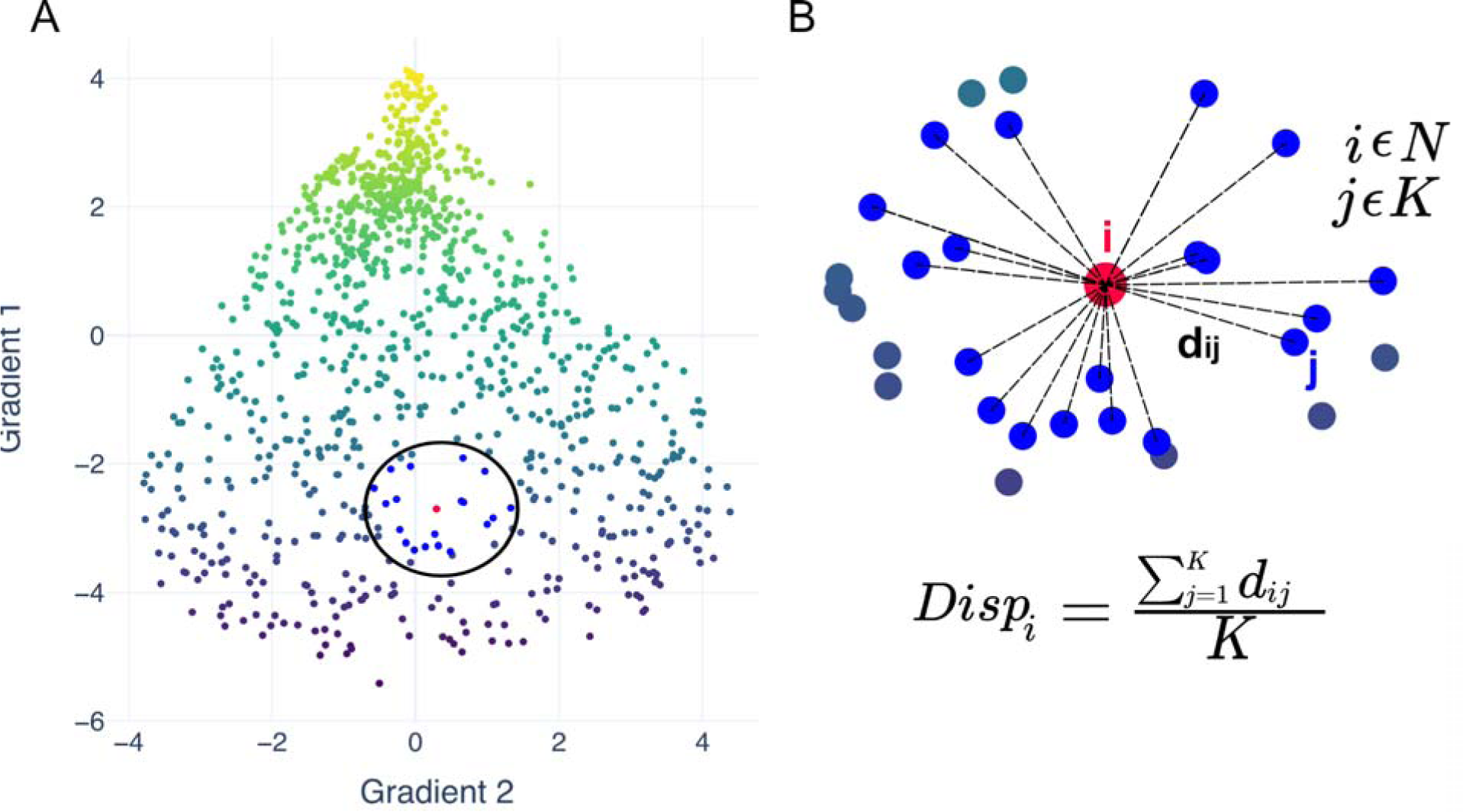
Illustration of neighborhood dispersion. **A:** In a multidimensional gradient embedding, for a given region (red) K closest neighbors are identified (blue). These regions are shown within the black circle. **B:** Gradient (neighborhood) dispersion of a given region is the mean distance between said region and its K closest neighbors. The same operation is done for every region (N regions = 1000).

Unlike previous investigations where the primary source of variability in dispersion originated from network delineation, our study has its own unique challenge in determining the number of the nearest neighbors (K). To address this issue, we included all dispersion values computed based on a range of nearest neighbors from 10 to 170 with a step size of 40. Basing the calculation of dispersion on differing sets of gradients (each including up to 200 gradients), we derived 1000 x 200 x 5 = 1,000,000 dispersion values, which we then used as input to our analytic workflow (along with flattened connectivity matrices and gradients).

### Analytic Workflow

The objective of our workflow was to identify the features with the largest predictive capacity from an extensive array of connectivity-based features. Our dataset included vectorized connectivity matrices (N = 499,500), 200 gradients (N = 200,000; 200 values for each region), neighborhood dispersion (N = 1,000,000) and centroid dispersion (N = 28). The schematic representation of the feature selection procedure is depicted in Figure 4. For each participant, we combined all features into one flat vector. Then, we concatenated the vectors for all 936 participants, resulting in a matrix of size [936 x 1,699,528] (Figure 4B, 1). Next, we applied principal component analysis (PCA) to each type of features separately and retained 20% of variance for each type (effectively compressing the feature dimension), except for the centroid dispersion for which all variance was included (Figure 4B, 2). The aim of the decomposition was to retain the same amount of variance for all feature types and to ensure that for each feature type more than one component is extracted when applying PCA. We retained 149 components in total (N_conn_ = 7, N_grad_ = 72, N_centroid_disp_ = 28, N_cortex_disp_ = 42; Figure 4B, 2.). The decomposed dataset was divided into the train (75% of participants) and holdout (25%) sets, with the ratio of N_SCZ_/N_NC_ = 0.35 in both.

**Figure 4.**
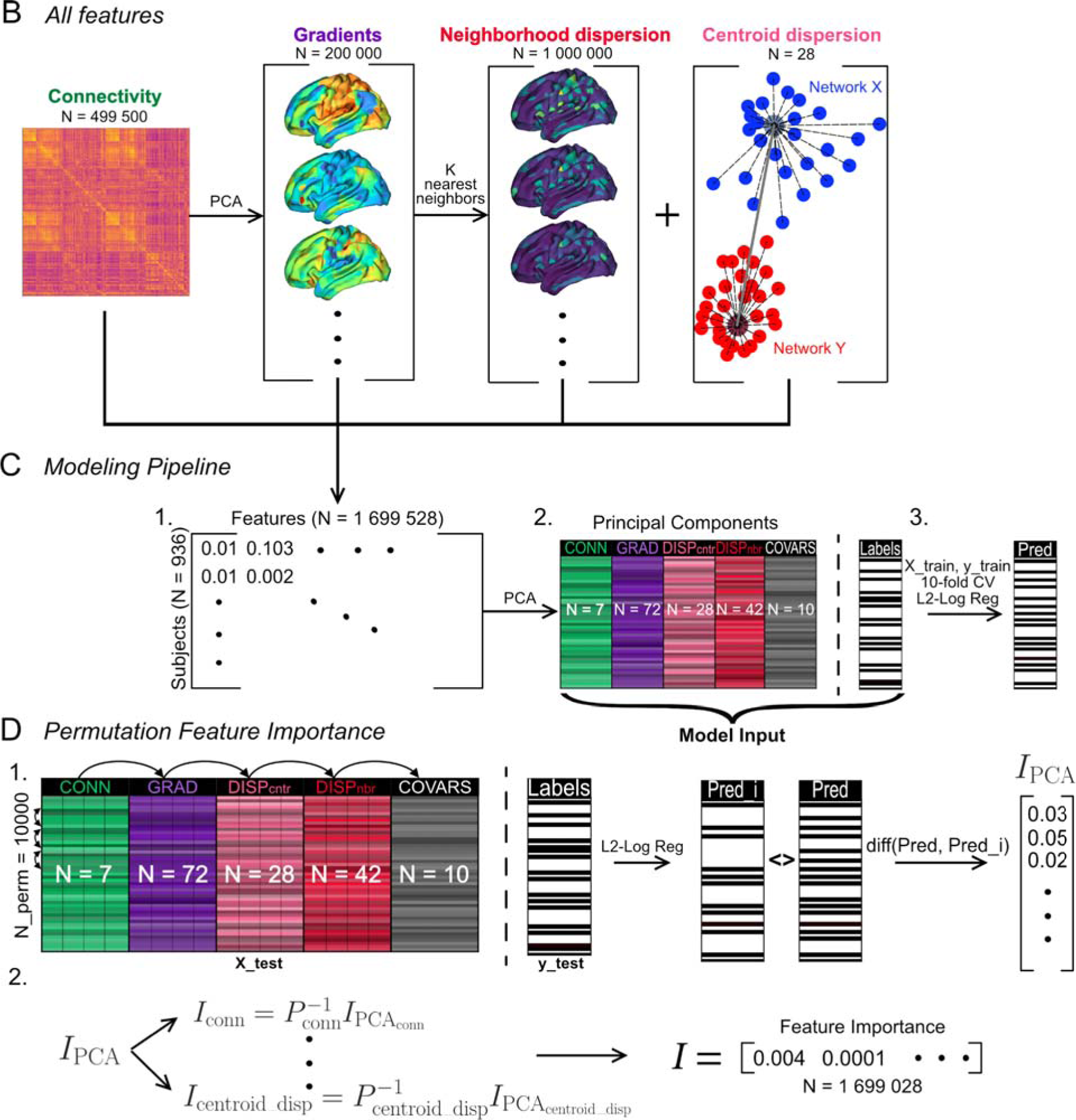
**A:** The types of predictors tested in this work (left to right): connectivity matrices (vectorized), macroscale cortical gradients, neighborhood, and centroid dispersion. **B:** All four types of features are concatenated together (2) and decomposed using group PCA (2) (each feature group is decomposed separately). The resulting dataset, along with covariates, was divided into the train and holdout set; 10-fold cross-validation (CV) was used to assess the performance of L2-regularized logistic regression on the PCA dataset (3). **C:** Permutation component importance was computed for each component using the holdout set (1). For each feature type, component importance was imverse transformed to obtain feature importance (2).

The purpose of the following steps was to quantify the importance of each component for classification performance. We fitted an L2-regularized logistic regression on the train set (Figure 4B, 3). Logistic regression was determined as the best model to compute component importance since the resulting coefficients are interpretable and the logic behind their computation is well understood. We used *permutation feature importance* (Figure 4C, 1) as the measure of the contribution of each component and feature to classification performance. Initially conceived for random forests^33,34^, it allows us to estimate the importance of the features in a classifier-agnostic way using the holdout set. First, the classifier is trained on the train set and the baseline accuracy on the holdout set is obtained. Second, each feature of the holdout set is randomly shuffled N_perm_ times and at each shuffle the permutation accuracy is computed. Permutation feature importance is the difference between the baseline accuracy and the permutation accuracy at each permutation. This importance metric can be estimated for any classifier; it quantifies the extent to which classification performance deteriorates or improves as every feature is shuffled. The feature with the largest permutation importance contributes the most to classification performance (Figure 4C, 1). We computed mean permutation importance for each component (N_perm_ = 10,000) and inverse transformed it based on the components’ projection matrices to obtain feature importance in the feature space (Figure 4C, 2). Permutation feature importance enabled the selection of features for further assessment of their utility for classification.

### Classifier Analysis

Finally, we assessed the predictive capacity of each feature type in a classifier-agnostic manner so as to prevent the results from being driven by the choice of a specific classifier. To this end, we selected 936 features with the largest permutation feature importance from each type and we trained and tested 13 distinct classifiers on them:

– Logistic regression (L2-regularized, LR).
– K-Neighbors Classifier (KN).
– Naïve Bayes (NB).
– Decision Tree Classifier (DT).
– Support Vector Machine (SVM).
– Ridge Classifier (Ridge).
– Random Forest Classifier (RF).
– Ada Boost Classifier (AB).
– Gradient Boosting Classifier (GB).
– Light Gradient Boosting Machine (LGB).
– Linear Discriminant Analysis (LDA).
– Extra Trees Classifier (ET).
– Quadratic Discriminant Analysis (QDA).

To verify that the difference in classification performance between feature types persists for larger numbers of features, we repeated this analysis for a range of 100 to 10,000 features with the largest permutation feature importance from each type.

For each feature count, we identified the best classifier based on its mean cross-validation (CV) accuracy across 10 folds. Next, we tested the best classifier on the holdout set. All data transformations were done within Scikit-Learn pipelines. The multi-classifier assessment was done using Pycaret (https://github.com/pycaret/pycaret). Age, sex, dataset of origin, and framewise displacement (FD) were always included as covariates.

For classifier performance, we report both accuracy and the F1-score. The F1-score, calculated using the Scikit-Learn package in Python, represents the harmonic mean of precision and recall. This metric is particularly useful in cases where class distribution is imbalanced, as it accounts for both false positives and false negatives.

The CV and test performance across all classifiers and feature subsets was compared to two baselines, namely:

1. Test performance on the PCA dataset (all 149 components) of the logistic regression.
2. the dummy classifier which randomly picks the class for each sample. It is frequently used as a baseline in machine learning research^35,36^.

Note that in this study the accuracy of the dummy classifier consistently remained at 73.7%. In contrast, the F1 score for this classifier is always 0, meaning that it does not differentiate between the two classes. These values constitute our chance reference.

## III. Results

### Permutation Feature Importance & Classification Performance

Upon visual inspection, we observed that the principal components of functional connectivity had the largest permutation feature importance, followed by gradients, centroid dispersion, and neighborhood dispersion (Figure 5A). We sought to verify that permutation feature importance indeed reflects an advantage in classification performance regardless of classifier. To this end, we selected the 936 features (as many as subjects) with the largest permutation feature importance for each type and trained and tested 13 classifiers on them (Figure 5). Connectivity outperformed the other feature types. This conclusion was supported by the Mann-Whitney U test (Figure 5B): when trained on connectivity, the classifiers had significantly higher F1-score than when trained on the 1^st^ (principal) gradient (U = 164, p = 0.003), neighborhood dispersion (U = 169, p = 0.001) and centroid dispersion (U = 156, p = 0.009). The other contrasts did not survive the correction for multiple comparisons (Bonferroni: α = 0.013).

**Figure 5.**
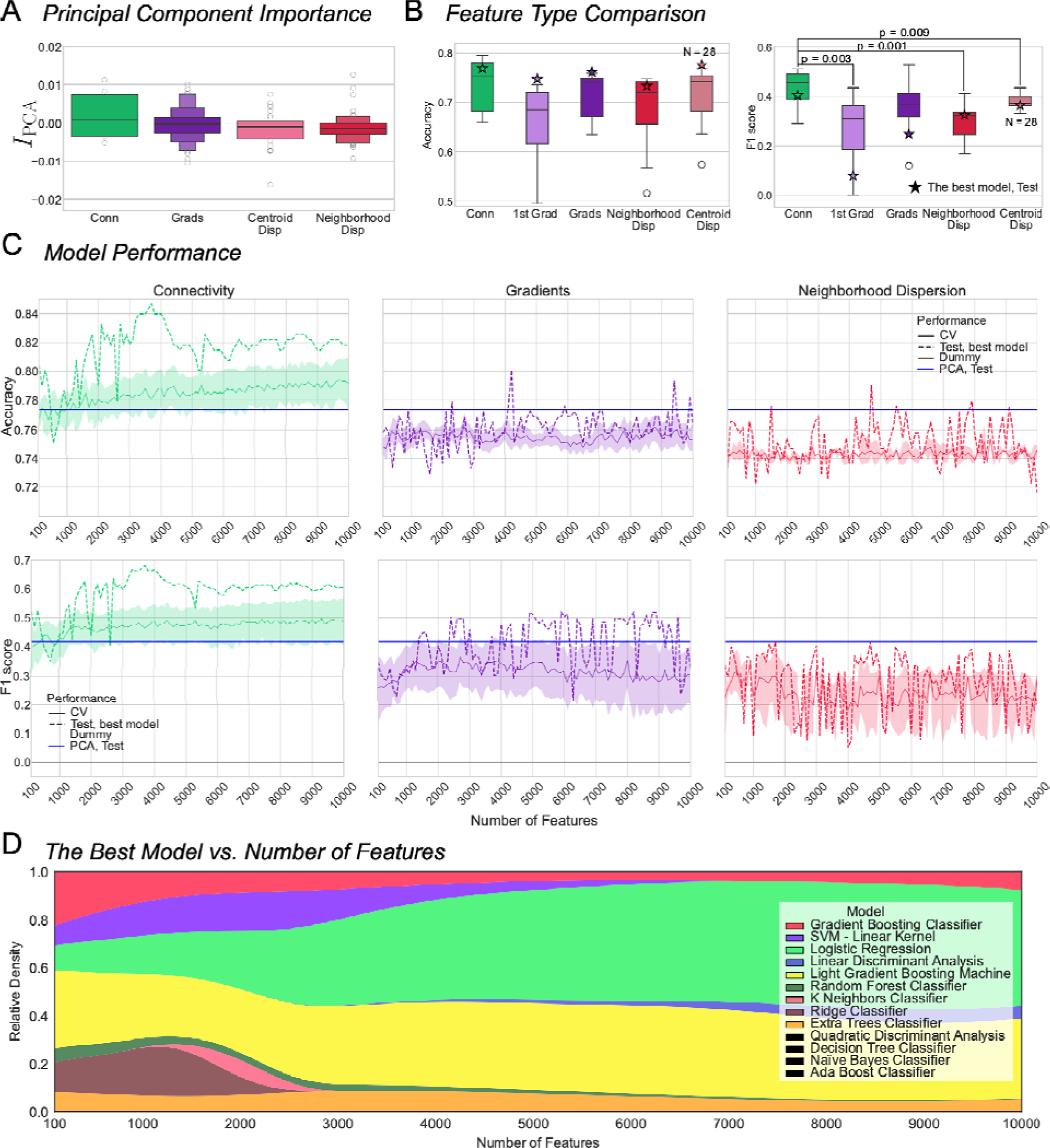
**A.** Permutation importance across feature types. **B:** Accuracy and F1 score across 13 classifiers (mean cross CV folds; the classifiers performing worse than the dummy classifier were excluded) fit on 936 best features from connectivity, the principal gradient, all gradients, neighborhood dispersion and the 28 values of centroid dispersion (light red). P-values indicate significant difference as per Mann-Whitney U test, α ≤ 0.013 (connectivity vs. all: Bonferroni-corrected). The stars denote the performance of the best classifier as identified based on the mean accuracy across 10 CV folds. **C:** Mean ± s.e.m. CV and test performance across classifiers for N features 100-10,000 for connectivity (left), gradients (middle), and neighborhood dispersion (right). Horizontal lines represent test performance of the logistic regression on all principal components (blue), and the performance of the dummy classifier (brown). The shading indicates s.e.m. **D:** Relative density of fits where the corresponding classifier was identified as best. Larger area indicates that the corresponding classifier had the highest CV accuracy more often. The legend features all classifiers that were tested in this study; the classifiers in black were never identified as the best.

Multi-classifier analyses further emphasized the superior predictive capacity of functional connectivity compared to the other feature types. Here we report CV and test accuracy and F1-score across all classifiers and for the best classifier (Figure 5C). Connectivity edges with the largest permutation feature importance consistently outperformed logistic regression fitted on all feature components. The other feature types performed substantially worse.

In addition, we examined the impact of the number of best features on the choice of the best classifier (fits on all feature types are considered). Figure 5D illustrates the evolution of the best classifier with the increasing number of features as the change in relative density of instances where the classifiers were identified as best. Three main patterns can be noted. Firstly, several classifiers clearly performed better with N_features < 3000: GB, SVM, RF, KN and Ridge. Secondly, ET and LGB had a relatively stable winning rate across all feature subsets. Thirdly, LR and LDA had an increased performance when N_features > 3000. However, LDA rarely outperformed the other classifiers, whereas LR and LGB often emerged as optimal.

### The Features with The Largest Permutation Feature Importance

To get a better understanding of the contribution of the most important connectivity features to the diagnosis prediction, we conducted an additional exploratory analysis. For three feature subsets including 500, 1000 and 5000 connectivity edges with the largest permutation feature importance, we computed weighted degree centrality. For each subset, the selected edges were transformed back to a 1000×1000 connectivity matrix and for each region the sum of edges was computed, each edge weighted by its correlation value. Thus, for each region, we sought to quantify its connectivity strength to the other regions given the selected edges.

For each edge subset, we plotted the difference in group degree centrality between SCZ and NC (Figure 6). Degree centrality was overall lower in patients with schizophrenia which is in line with previous accounts of lower overall functional connectivity characteristic of this disease^37–42^. This finding lends support to the dysconnectivity hypothesis^43,44^.

**Figure 6.**
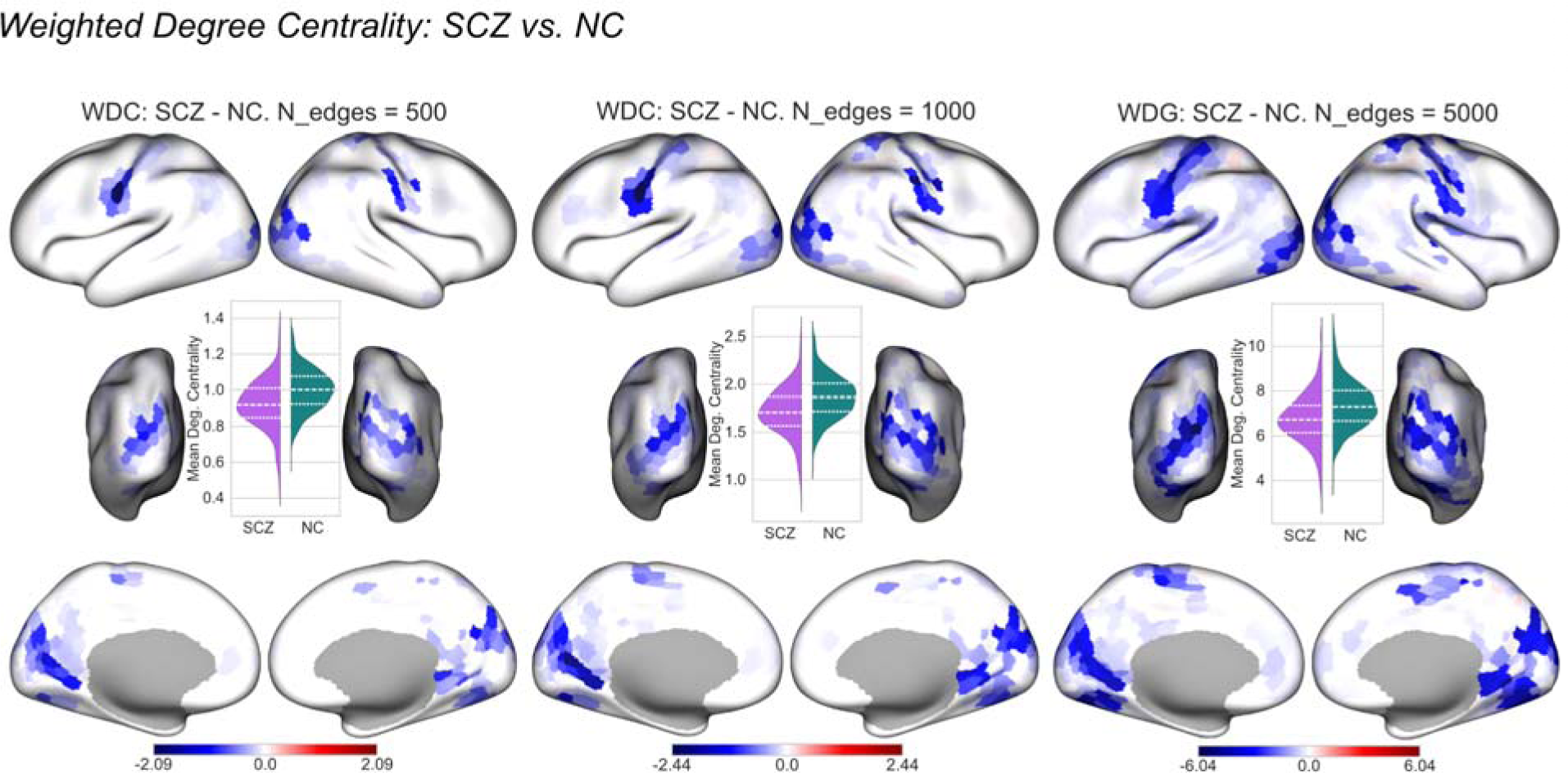
The difference in weighted degree centrality between the two groups for 500, 1000 and 5000 connectivity edges with the largest permutation feature importance. Inset violinplots display weighted degree centrality averaged across regions for the two groups. SCZ: patients with schizophrenia, NC: neurotypical controls, WDG: weighted degree centrality.

Secondly, the spatial pattern illustrated in Figure 6 indicated that the significant differences in degree centrality were concentrated in the primary regions for all edge subsets displayed here. Put differently, the edges with the largest permutation feature importance reflect connectivity strength in the primary regions, indicating that their connectivity profiles contribute the most to classification performance.

## IV. Discussion

Here, we extend the effort to determine the optimal connectivity-based predictors of behavior (R. Kong et al., 2023) to psychopathology by attempting to benchmark connectivity-based features against each other for the prediction of the diagnosis of schizophrenia. To this end, we applied a feature selection workflow based on permutation feature importance to a large dataset comprising connectivity, macroscale cortical gradients and gradient dispersion. Our analysis revealed that, despite growing interest in cortical gradients and gradient dispersion^12,15,18,45^, functional connectivity holds superior predictive potential for schizophrenia over its low-dimensional derivatives (Figure 5B, C). Additionally, we demonstrated that the connectivity edges connecting the primary sensory regions have the largest permutation feature importance. This result indicates that variations in connectivity strength of these regions encapsulate critical information for distinguishing patients from controls. This finding, however, does not discard the relevance of the gradients for the studies looking specifically into functional hierarchical variations. Regarding our use case, the diminished predictive capacity of the gradients could stem from the conservative matrix thresholding applied prior to dimensionality reduction. Future studies are needed to test this hypothesis.

To gain a deeper understanding of the connectivity patterns associated with schizophrenia, we conducted an exploratory analysis whereby we compared weighted degree centrality for the regions linked by the 500, 1000 and 5000 most important edges. Firstly, degree centrality appeared to be lower in patients with schizophrenia, corroborating previous evidence of overall hypoconnectivity typical for the disorder^37–42^. Secondly, for the edges with the largest importance, these differences were predominantly concentrated in the sensorimotor, auditory, and visual cortex. These findings appear to contrast with the studies highlighting discrepancies predominantly in higher-order areas, such as the default mode network (DMN)^20,46–49^. However, the differences we observed need to be contextualized as the ones relevant for classification performance: while they may or may not constitute the neurological basis of schizophrenia, they are most informative for the differentiation of the two groups. Viewed from this perspective, our results resonate with the research demonstrating that across individuals the measurements are most reliable in the primary, unimodal areas^50,51^. In addition, one study reported more accurate surface registration for the primary sensory areas^52^. It has been postulated that this stability can be attributed to the fact that the primary areas are phylogenetically the most ancient cortical areas and are therefore considered as evolutionary anchors around which most of the cortical expansion in humans unfolded^31,51,53–55^. The largest feature importance in the primary and most stable regions may indicate that the classifiers rely on those individual differences which are replicable across individuals of the same group. In such a setting, intra-individual variability cannot be accounted for. This shortcoming can be potentially addressed by studies focusing on datasets featuring hours of scanning time per individual^56–58^.

We were also interested in how the ranking of the 13 classifiers tested in this work evolved with the number of best features selected based on permutation feature importance. As depicted in Figure 5D, some classifiers’ performance relative to each other fluctuated considerably depending on the size of the feature subset. Overall, no classifier outperformed others at all times for any number of features. It is a clear indication of the necessity of empirical testing of the candidate classifiers. However, it appears that as the number of features increases, the performance of simpler classifiers — such as logistic regression and LDA — improves.

Regarding limitations, a certain degree of caution is warranted when interpreting our results. Firstly, we did not account for the effect of medication in our study since the medication data were missing for a large number of patients. Excluding these subjects would have resulted in a drastically decreased sample size and, more importantly, in an increased imbalance of classes.

Secondly, most classifiers in this study were fitted on datasets with more features than observations which increases the risk of overfitting. Nonetheless, we believe to have addressed this issue by employing a 10-fold CV for each classifier. In addition, we have also witnessed that for some classifiers test performance exceeded CV performance (Figure 5C) which is not characteristic of overfitting.

In addition, we did not consider the issue of comorbidity and transdiagnostic phenomena across psychiatric disorders. While this topic lies beyond the purview of the present study, it is imperative to acknowledge its critical significance. The inquiry into transdiagnostic classification and symptom prediction holds potential to profoundly transform the paradigms governing the diagnosis and treatment of patients. Indeed, as of late, the research looking into transdiagnostic effects has garnered substantial momentum^5,59–64^.

The emergence of novel connectivity-based methods broadens our toolkit for predicting psychiatric disorders, introducing a necessity for empirical validation. Our findings indicate that functional connectivity outperforms its more recent, low-dimensional derivatives such as cortical gradients and gradient dispersion in predicting schizophrenia. Additionally, in this study, the connectivity within the primary sensory regions showed the highest discrimination capabilities, possibly due to the reduced anatomical and functional variability of those regions. We anticipate, however, that it is also informative for a broad spectrum of major psychiatric disorders. The exploration of this latter possibility warrants thorough examination in future work.

## Acknowledgements

We thank Joshua Vogelstein for methodological guidance and Vincent Kreft for contributing to the conceptual development of the study. This project has received funding from the European Research Council (ERC) under the European Union’s Horizon 2020 research and innovation programme (Grant agreement No. 866533) awarded to D.S.M. This project was also supported by the *contrat doctoral spécifique normalien (CDSN)* awarded to V.S. by the Ministry of Higher Education and Research of France. The computational aspects of this research were supported by the Wellcome Trust Core Award Grant Number 203141/Z/16/Z and the NIHR Oxford BRC. The views expressed are those of the author(s) and not necessarily those of the NHS, the NIHR or the Department of Health.

## Author contributions

S. researched the data, conducted the analyses and wrote the manuscript. C.P., D.W., P-L.B., and D.S.M provided methodological expertise. C.P., R.A.B, R.S., W.W., U.K., F.A., D.W., T.D.S., P-L.B., and D.S.M. assisted in interpretation of findings and manuscript revision.

## Data availability

The data used in this work are in open access, conditioned upon the signature of a data sharing agreement; for COBRE: http://schizconnect.org/, SPBRS-1600: https://bicr-resource.atr.jp/srpbsopen/, and LA5c: https://openfmri.org/dataset/ds000030/.

## Computer code

The code used to perform the analyses and to produce the figures featured in this study is available Github: https://github.com/victoris93/feature-selection-scz.

