## Supplementary Methods for "A Comparative Machine Learning Study of Connectivity-Based Biomarkers of Schizophrenia"

### Supplementary Table 1: Scanning Parameters

| Dataset | COBRE | LA5c | SWA[1] | KUT[1] | KTT[1] | UTO[1] |
|---|---|---|---|---|---|---|
| Site | UNM Center for Psychiatric Research | UCLA | Showa university | Kyoto university | Kyoto university | University of Tokyo |
| MRI Scanner | SIEMENS TrioTim | SIEMENS TrioTim | SIEMENS Verio | SIEMENS TrioTim | SIEMENS Trio | GE Discovery MR750w |
| Magnitic field | 3T | 3T | 3T | 3T | 3T | 3T |
| TR (s) | 2 | 2 | 2.5 | 2.5 | 2 | 2.5 |
| TE (ms) | 29 | 30 | 30 | 30 | 30 | 30 |
| Flip angle (deg) | 75 | 90 | 80 | 80 | 90 | 80 |
| Field of view (mm) | 256 | 192 | 212 | 212 x 212 | 256 x192 | 212 |
| In-plane resolution (mm) | 3.75 × 3.75 | NA | 3.3 x 3.3 | 3.3125 x 3.3125 | 4.0 x 4.0 | 3.3 |
| Slice thickness (mm) | 3.5 | 4 | 3.2 | 3.2 | 4.0 | 3.2 |
| Slice gap (mm) | 4.55 | NA | 0.8 | 0.8 | 0 | 0.8 |
| Number of slices | 33 | 34 | 40 | 40 | 30 | 40 |
| Total scan time | 6:00 | 5:12 | 10:17 | 10:10 | 06:00 | 10:10 |

### Supplementary Methods 1: MRI Preprocessing, fMRIPrep

The following preprocessing steps were applied to each subject. A reference volume and its skull-stripped version were generated using a custom methodology of fMRIPrep. Susceptibility distortion correction (SDC) was omitted. The BOLD reference was then co-registered to the T1w reference using flirt (FSL 5.0.9[2]) with the boundary-based registration[3] cost-function. Co-registration was configured with nine degrees of freedom to account for distortions remaining in the BOLD reference. Head-motion parameters with respect to the BOLD reference (transformation matrices, and six corresponding rotation and translation parameters) were estimated before any spatiotemporal filtering using mcflirt (FSL 5.0.9[2]). The scans were slice-time corrected using 3dTshift from AFNI 20160207[4] (RRID:SCR_005927). The BOLD time-series (including slice-timing

[1] These datasets are part of a larger dataset, SPRBS-1600[1].

correction) were resampled onto their original, native space by applying the transforms to correct for head-motion. These resampled BOLD time-series will be referred to as preprocessed BOLD.

The BOLD time-series were resampled into standard space, generating preprocessed BOLD in MNI152NLin2009cAsym space. First, a reference volume and its skull-stripped version were generated using a custom methodology of fMRIPrep. Several confounding time-series were calculated based on the preprocessed BOLD: framewise displacement (FD), DVARS and three region-wise global signals. FD was computed using two formulations following Power (absolute sum of relative motions)[5] and Jenkinson (relative root mean square displacement between affines)[2]. FD and DVARS were calculated for each scan using their implementations in Nipype (following the definitions by Power et al.[5]). The three global signals were extracted within the cerebrospinal fluid (CSF), the white-matter (WM), and the whole-brain masks. The frames that exceeded a threshold of 0.5 mm FD or 1.5 standardised DVARS were annotated as motion outliers. We regressed out the following artifacts: CSF, WM, and head motion confound timeseries and their quadratic derivatives. The full list of confound regressors as output by fMRIprep:

```
csf
white_matter
trans_x
trans_x_derivative1
trans_x_derivative1_power2
trans_x_power2
trans_y
trans_y_derivative1
trans_y_power2
trans_y_derivative1_power2
trans_z
trans_z_derivative1
trans_z_power2
trans_z_derivative1_power2
rot_x
rot_x_derivative1
rot_x_derivative1_power2
rot_x_power2
rot_y
rot_y_derivative1
rot_y_derivative1_power2
rot_y_power2
rot_z
rot_z_derivative1
rot_z_power2
rot_z_derivative1_power2
```

After initial preprocessing with fMRIprep, we further preprocessed the signal using Nilearn (Python). The timeseries were corrected for temporal drifts and trends and normalized (z-score). Finally, low- and high-pass filtering was applied at 0.01-0.08 Hz.
